# Variations in δ^13^C of different plant organs: implications for post-photosynthetic fractionation

**DOI:** 10.1101/238477

**Authors:** Yonge Zhang, Xinxiao Yu, Lihua Chen, Guodong Jia

## Abstract

Compared to photosynthetic fractionation, the mechanism of post-photosynthetic carbon isotope fractionation is not well understood. The aim of this study was to investigate post-photosynthetic fractionation in both above and below ground tissues and to evaluate potential hypotheses explaining differences in carbon isotope composition (δ^13^C) among different plant organs, which can provide valuable insights into plant physiology. The results revealed that there is no significant day-night difference in *δ*^13^C of twig phloem water soluble organic materials (WSOM), which could be explained by the unrestricted exchange of triose-phosphates between the chloroplast and cytoplasm and a time lag for carbohydrate exportation. Further, we found that *δ*^13^C of twig phloem WSOM is more sensitive to plant water status than leaf WSOM. Analysis of *δ*^13^C in different plant organs showed that the greatest ^13^C enrichment was recorded in stem phloem. Divergences in *δ*^13^C of phloem WSOM among different plant organs were not likely to be explained by respiratory fractionation or time lag and were ascribed to transport of carbohydrates across organ boundaries and metabolic processes. Our study demonstrated that post-photosynthesis fractionation could not be ascribed to a single, unifying hypotheses; instead, it is the result of multiple processes.

**Highlight:** *δ*^13^C of twig phloem water soluble organic materials varied no clear diel pattern. In the leaf-twig-stem-root sequence, the greatest ^13^C enrichment was recorded in stem phloem.

## Introduction

^13^C fractionation during photosynthesis had been shown to produce divergence in carbon isotope composition (*δ*^13^C) between the atmosphere and plant organic material, leading to leaves that are ^13^C enriched in comparison to the atmosphere (Seibt *et al*., 2008; Hu *et al*., 2010). After sugars are synthesized in the chloroplast, some of the new assimilates are then exported from leaves and transported in the phloem to different plant organs, such as twigs, stems, and roots (Mahboubi, 2014). To date, a large number of studies have reported that *δ*^13^C of heterotrophic organs can be altered, resulting in distinct signatures in different plant organs because of carbon isotope fractionation following carboxylation (Badeck *et al*., 2005; Eglin *et al*., 2009; Zhao *et al*., 2017). Post-photosynthetic carbon isotope fractionation has been associated with activities such as transport of metabolites, the process of respiration, and biosynthesis of organic materials (Damesin and Lelarge, 2003; Gessler *et al*., 2004; Brandes *et al*., 2006; Bathellier *et al*. 2008). In contrast to photosynthetic carbon isotope fractionation, the mechanism of post-photosynthetic fractionation in different plant organs is not clear. Cernusak *et al*. (2009) proposed six hypotheses that may explain the divergence in *δ*^13^C between source and sink tissues. In addition, Badeck *et al*. (2005) reviewed more than 80 studies and found that autotrophic tissues were generally ^13^C-depleted in comparison to heterotrophic tissues. However, there are no definitive conclusions about carbon allocation among different organs owing to post-photosynthetic fractionation. Recently, post-photosynthetic fractionation has been extensively investigated, providing important insights into plant physiology. Further, the interpretation of post-photosynthetic fractionation allows the reconstruction of individual sinks and sources using δ^13^C of CO2 exchange flux at the ecosystem scale because δ^13^C of respired CO_2_ of different plant organs is closely related to that of the respiratory substrate (Bowling *et al*., 2001; Gessler *et al*., 2008; Wehr *et al*., 2015).

Considerable uncertainty and complexity, including spatial variation, complicates investigation of post-photosynthetic fractionation of tall adult trees with heterogeneous vertical structure under field conditions. To avoid spatial inconsistencies, we increased sample size and sampled leaf and twig material at different canopy heights. Most recent studies have interpreted post-photosynthetic fractionation based on total organic materials of both autotrophic and heterotrophic tissues (Badeck *et al*., 2005; Cernusak *et al*., 2009); however, this method is not very efficient when considering the process of carbon transport among different plant organs. Carbon isotopes from the fast-turnover carbohydrate pool (water-soluble organic materials, WSOM) in leaves are commonly used indicators of assimilation-weighted average *C*_i_/*C*_a_ (the ratio of intercellular to atmospheric CO_2_ concentration) integrated over a period of several hours to 1–2 days (Pons *et al*. 2009), which can be imprinted ininfluence phloem WSOM rapidly. Therefore, analysis of *δ*^13^C of WSOM can provide valuable information about post-photosynthetic fractionation within plants. Recently, the development of phloem collection techniques has enabled ^13^C signals of heterotrophic phloem WSOM to be precisely determined. In this study, we detected *δ*^13^C of WSOM for both above and below ground organs, which may contribute to understanding the carbon exchange process between above and below ground organs and the mechanism of interaction between plants and the soil (Deng *et al*., 2017).

The objectives of the study were to investigate post-photosynthetic fractionation in the leaf-twig-stem-root sequence, as carbohydrate is exported from leaves and transported in the phloem throughout the whole plant, and to evaluate the potential hypotheses that can explain divergence in *δ*^13^C among different plant organs.

## Materials and methods

### Study site description

The research was conducted at the Jiufeng National Forest experimental station, located in Beijing in a rocky, mountainous area of north China (116°05’E, N40°03’N). The climate is a typical warm temperate, semi-humid, and semi-arid monsoon climate. Annual average temperature and precipitation are 11.6 °C and 650 mm, respectively, and average evaporation ranges from 1,800 to 2,000 mm per year. Precipitation occurs largely between July and September, which is much less than evaporation. The growing season in this area lasts from April to October. The soil is sandy loam soil with mean thickness of about 52.8 cm. Drought-tolerant conifers are the main tree species.

The central area (about 1 km^2^) of the sampling site mainly consists of 50–60-year-old *Platycladus orientalis*, with an average tree height and breast diameter of 8 m and 20 cm, respectively, and forest crown density is about 0.8. There are small patches of shrubs and herbs, predominantly *Vitex negudo, Grewia biloba*, and *Broussonetia papyrifera*.

### Sampling and *δ*^13^C determination of different plant organs

Sampling was performed on several sunny days in 2017. On 25 May, 17 July, and 29 August, samples were collected every two hours from 1:00 to 24:00 for analysis of diel variation in carbon isotopes of different plant organs. In addition, on 19 April, 8 August, 10 August, and 11 August, plant materials were collected every six hours from 1:00 to 24:00 to calculate daily mean values. On each sampling day, three mature *Platycladus orientalis* conifer leaves and small twigs were collected in each canopy crown (including the upper, middle, and lower canopy), and three replicate stem phloem and fine root samples were also collected. After sampling, all plant materials were covered with foil and rapidly preserved in liquid nitrogen. All samples were transported to the Key Laboratory of Soil and Water Conservation and Desertification Combating of Ministry of Education of Beijing Forestry University for carbon isotope composition (δ^13^C) analysis.

To obtain leaf water-soluble organic material (WSOM), the middle veins of the leaves were removed. Next, 50 mg of freshly abraded leaf samples were well mixed with 100 mg PVPP (polyvinylpyrrolidone) and double demineralized water up to 1.75 mL was added in a 2.0 mL centrifuge tube. These mixtures were incubated for 1 h at a constant temperature of 5 °a. The small twig, stem, and root phloem WSOM were collected according to the methods of Brandes *et al*. (2006). Similar to the collection of leaf WSOM, 75 mg phloem samples were incubated in 1.75 mL double demineralized water in a 2.0 mL centrifuge tube at room temperature for 24 h. Subsequently, the leaf and phloem materials were boiled at 100 °C for 3 min and centrifuged for 5 min (12000 g). Supernatants of WSOM (10 μL) were transferred into tin capsules and oven-dried at 70 °C before *δ*^13^C analysis. Finally, *δ*^13^C of dry residues were analyzed using a stable isotope ratio mass spectrometer (Finnigan DELTA^plus^XP, Thermo Fisher Scientific, USA).

### Meteorological measurements

A meteorological tower at the sampling site continuously output micrometeorological data. Air temperature (*T*_a_) and relative humidity *(RH)* sensors (HOBO-U30, Onset, USA) were installed in the lower (5 m), middle (6.5 m), and upper crown (8 m). Vapor pressure deficit *(VPD)* was calculated as *VPD*=0.611×10^[17.502*T*a(240.97+*T*a)]^× (1-RH).

Additionally, solar radiation (*R*_a_) was measured at 8 m by a HOBO-U30 sensor (Onset, USA). Further, soil water content (*M*_s_) was determined using three sensors (HOBO-U30, Onset, USA) which were installed 10–30 cm below the ground. All micrometeorological data were recorded every 10 min.

## Results

### Measurements of meteorological conditions

Both *T*_a_ and *VPD* exhibited similar diel dome-shaped variation at each sampling period, reaching maximum values around noon and minimum values during the night (Fig. 1). However, *RH* had the opposite trend over time. Among canopy heights, maximum *T*_a_ and *VPD* were generally recorded in the upper canopy and minimum values were typically observed in the lower canopy. In contrast, *RH* tended to decrease with increasing canopy height (Fig. 1).

**Figure 1.**
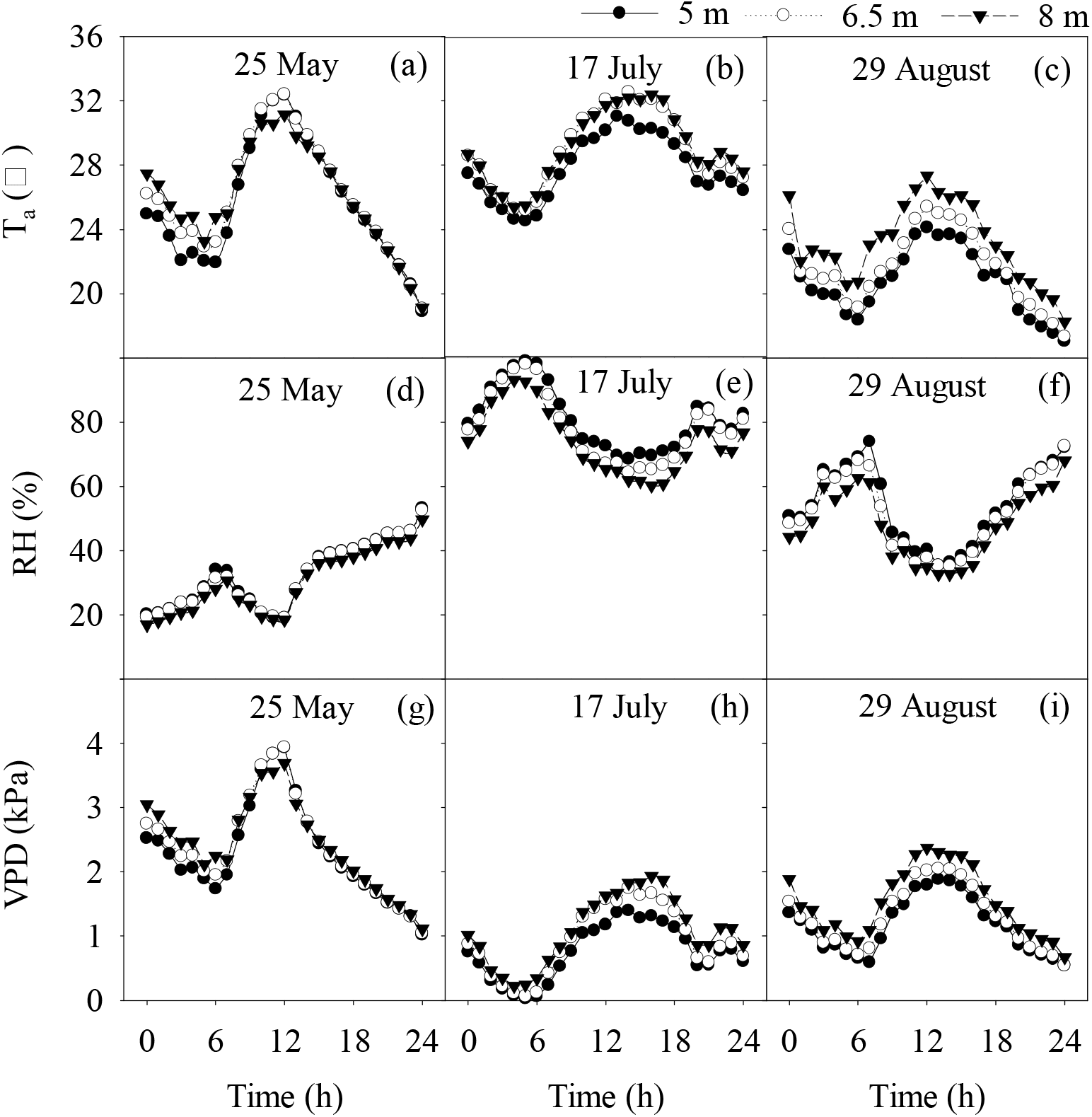
Diel variation in temperature (*T_a_*), relative humidity (*RH*) and vapor pressure deficit *(VPD)* on three measurement days at different canopy heights.

*M*_s_ was almost constant over diel time scales within the three sampling days, but it varied among the days (Fig. 2). *M*_s_ in the rainy season (on 17 July and 29 August) was significantly higher than that in the dry season. *R*_a_ had a varying diel dome-shaped pattern, reaching maximum values around noon, then decreasing to its minimum value in the evening (Fig. 2).

**Figure 2.**
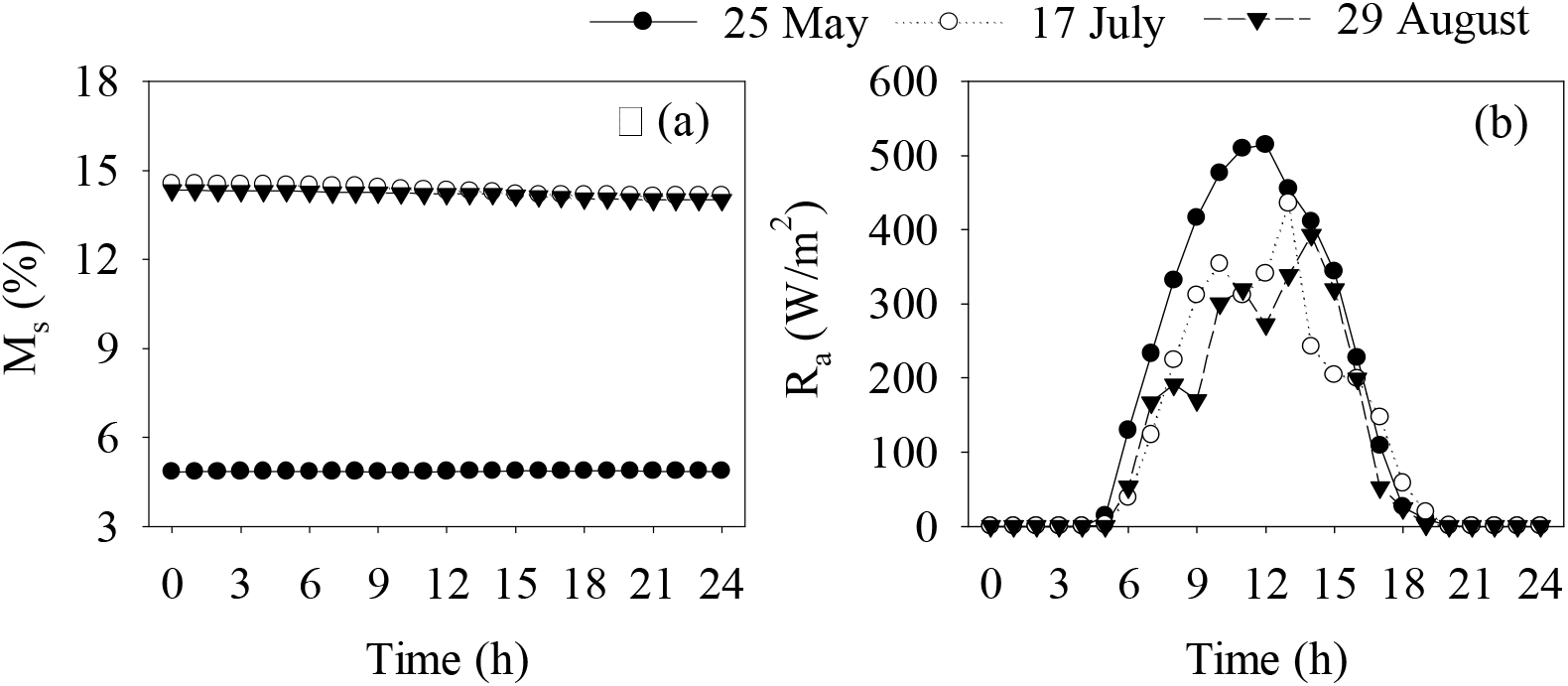
Diel variation in soil moisture (*M*_s_) and solar radiation (*R*_a_) on three measurement days at different canopy heights.

### *δ*^13^C of different plant organs

The *δ*^13^C of leaf and small twig phloem WSOM had no distinct diel pattern and ranged from −27.82‰ to −26.32‰ and from −28.18‰ to −26.47‰, respectively (Fig. 3). However, at different canopy heights, we repeatedly observed clear intracanopy gradients of *δ*^13^C of leaf and twig phloem WSOM. The greatest *δ*^13^C enrichment of leaf and twig phloem WSOM was observed in the upper canopy, and *δ*^13^C in the lower canopy was more negative in comparison with those in the middle and upper canopy.

**Figure 3.**
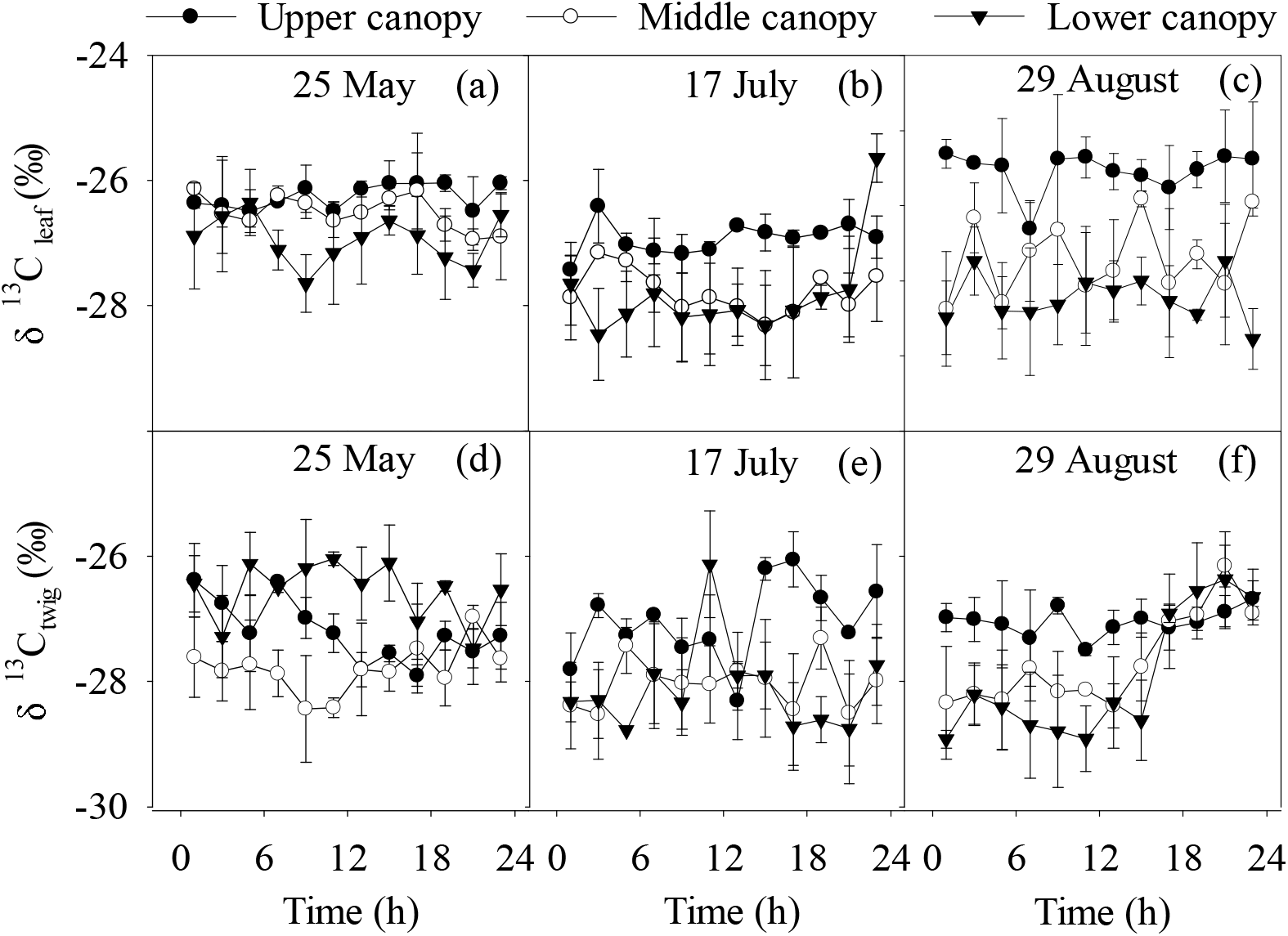
Intracanopy gradients of *δ*^13^C of leaf and twig phloem WSOM. Data are mean values ± SD.

The *δ*^13^C analysis of both autotrophic tissue (leaves) and heterotrophic tissue phloem (small twigs, stems, and roots) WSOM is shown in Fig. 4. Although there were no significant day-night differences in *δ*^13^C of these plant organs (*p*>0.05), there were clear differences among different plant organs. The greatest ^13^C enrichment was observed in stem exudates, which ranged between −26.21‰ and −24.48‰. *δ*^13^C of leaf WSOM was generally more positive than that of small twig phloem WSOM, with a mean difference of 0.44±0.36‰. On 25 May and 29 August, *δ*^13^C of root phloem WSOM was similar to that of leaf WSOM. However, on 17 July, root phloem WSOM was slightly enriched in *δ*^13^C compared to leaf WSOM.

**Figure 4.**
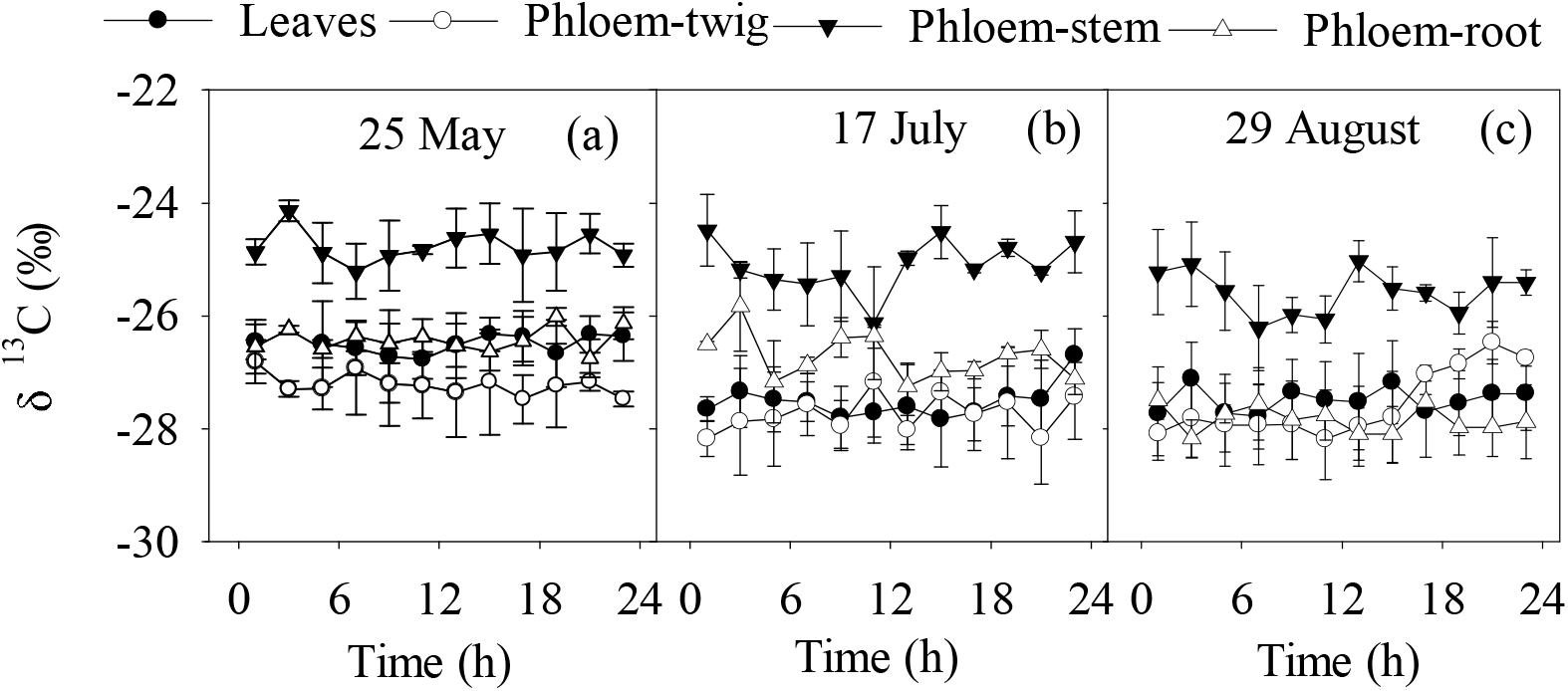
*δ*^13^C of WSOM of different plant organs; *δ*^13^C of leaf and twig phloem WSOM are means of the upper, middle, and lower canopy values. Data are mean values ± SD.

### Daily average values

Daily average values were analysed to describe day-to-day variation in *δ*^13^C of different plant organs. Fig. 5 shows that *δ*^13^C of WSOM of all plant organs had the most positive values on 25 May, and the second-highest values were recorded on 19 April, but the differences were not significant. The day-to-day variation of *δ*^13^C of small twig and stem phloem WSOM was similar to that of leaf WSOM. Further, daily mean *δ*^13^C values of root phloem WSOM tended to be more variable than those of WSOM of other plant organs.

**Figure 5.**
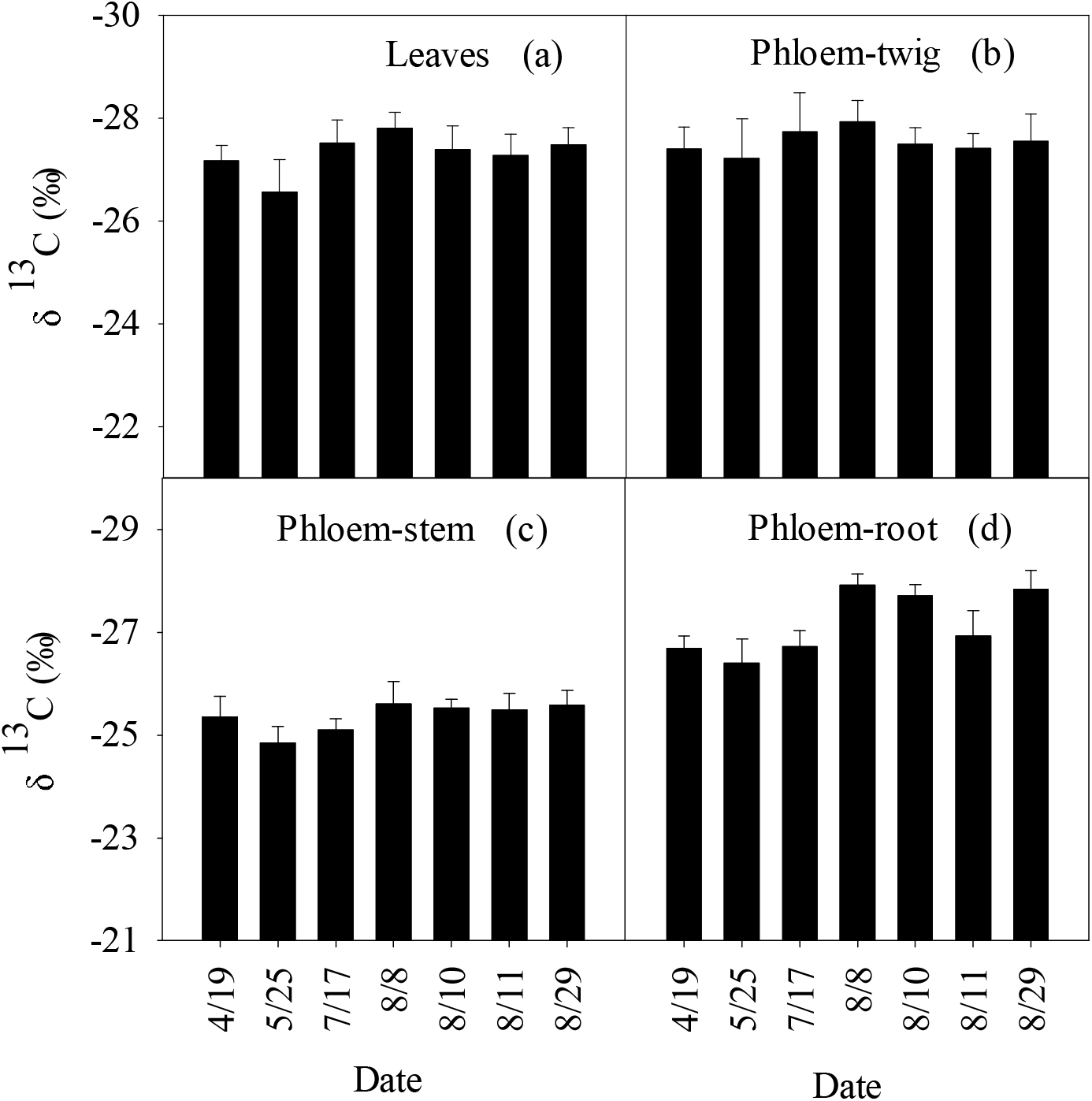
Day-to-day variation of *δ*^13^C of different plant organs collected on 19 April, 25 May, 17 July, 8 August, 10 August, 11 August, and 29 August. Error bars are the standard error of average values

### Correlation analysis

At smaller temporal scales (two-hour intervals), there were moderate correlations between *δ*^13^C of leaf WSOM and *RH* or *VPD*, with *r* of −0.62 and 0.63, respectively (*p*<0.01) (Table 1). The results showed that neither *δ*^13^C of WSOM of leaves nor that of twig phloem had significant correlations with *M*_s_. At larger temporal scales (daily mean values), there were significant correlations between *M*_s_ and *δ*^13^C of leaf or twig WSOM (*r* equal to −0.83 and −0.91, respectively, *p*<0.01), but the correlations between *RH* or *VPD* and *δ*^13^C of leaf or twig WSOM were not significant (Table 2).

**Table 1.** Correlation between environmental factors and *δ*^13^C of WSOM of different plant organs. (*n*=12, values are obtained at two-hour intervals at three sampling dates: 25 May, 17 July, and 29 August).

|  | Leaves | Twig-phloem | Stem-phloem | Root-phloem |
| --- | --- | --- | --- | --- |
| $T_a$ | -0.21 | -0.06 | 0.47** | 0.66** |
| $RH$ | -0.63** | -0.33 | -0.15 | -0.06 |
| $VPD$ | 0.633** | 0.26 | 0.35 | 0.30 |
| $M_s$ | -0.56** | -0.55* | -0.44* | -0.53** |
| $R_a$ | 0.31 | 0.02 | 0.17 | 0.24 |

\* $p<0.05$ ; \*\* $p<0.01$ .

**Table 2.**
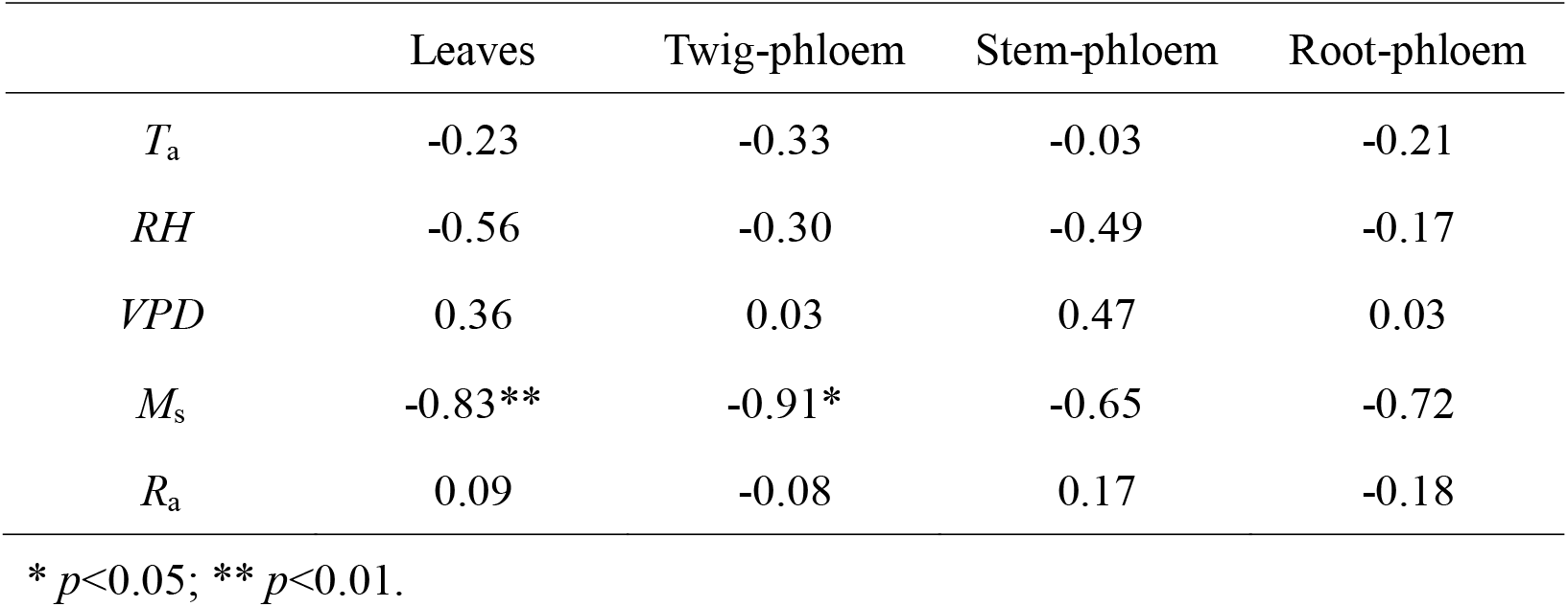
Correlation between environmental factors and *δ*^13^C of WSOM of different plant organs. (*n*=7, daily mean values).

## Discussion

### ^13^C enrichment of above ground organs

Previous studies have demonstrated that chloroplasts are the place where carbon assimilation occurs (Seibt *et al*., 2008; Bogelein *et al*., 2012). During light periods, large amounts of glucose are synthesized in chloroplasts as a result of their high photosynthetic capacity. Some of this glucose is exported to the cytosol by triose phosphate translocators (TPTs) for sucrose biosynthesis and other metabolic activities, and excess glucose accumulates and converts into starch in the chloroplast. In contrast, during dark periods, photosynthesis is constrained without illumination, and starch in the chloroplast is remobilized to produce glucose and maltose, which are exported to the cytosol to be utilized in the biosynthesis of sucrose and the process of primary metabolism (Cho *et al*., 2011; Mahboubi, 2014). During the course of aldolase-catalysed reactions in the chloroplast, the aldolase isotope effect bring about relatively ^13^C enriched hexoses will be incorporated in starch while lighter trioses will be exported to the cytoplasm. Therefore, *δ*^13^C in the chloroplast starch is more positive than that in the cytosolic WSOM in the daytime (Brandes *et al*., 2006; Cernusak *et al*., 2009). Instead, the breakdown of starch leads to sucrose being exported from leaves, which can carry the signature of the starch substrate. Previous studies determined that sucrose is the major carbohydrates that can be exported into phloem and transported within the whole plant (Gessler *et al*., 2008; Mahboubi, 2014). Peuke *et al*. (2001), observed that about 90% of the carbohydrates transported in the phloem is made up of sucrose. This can result in distinct day-night differences in δ^13^C of leaf and small twig phloem WSOM. In this study, *δ*^13^C analysis of diel variation revealed that there are no significant day-night differences in *δ*^13^C of the fast-turn-over carbon pool in leaves and in twig phloem (Fig.3). This finding is in contrast to previous results showing that *δ*^13^C in water-soluble leaf and twig compounds are generally enriched during night or early morning and relatively depleted during light periods. Our results may differ from these previous findings because the exchange mechanism of triose-phosphates between the chloroplast and cytoplasm may be species-specific. Tcherkez *et al*. (2004) proposed that aldolase-catalysed activities in the cytoplasm and chloroplast would utilize the same pool of triose-phosphates as substrates if the exchange was not restricted. In that case, δ^13^C of starch is quite similar to that of light-produced sucrose. Moreover, twig phloem WSOM is a mixture of carbohydrates, with different metabolic histories, derived from carbon exported from source tissues (leaves) during both light and dark periods. According to Cernusak *et al*. (2009), if leaves and small twigs were sampled at the same time during daylight, twig phloem could carry ^13^C signals originating from both day and night exports from source tissues (leaves). For example, Ruehr *et al*. (2009) suggested that it takes less than 5 hours for carbohydrate exportation when soil water content is high, while exportation increased to 4 days under drought conditions. As a consequence, day-night differences in *δ*^13^C of twig phloem WSOM are strongly inhibited.

Although diel variation in *δ*^13^C of WSOM of above ground organs was not clear, intracanopy gradients of *δ*^13^C of leaf and twig phloem WSOM were evident (Fig.3). Similar intracanopy gradients have been found in Douglas-fir leaves and twig phloem *δ*^13^C (Bogelein *et al*., 2012). This finding indicated that *δ*^13^C of leaf and twig phloem WSOM can be influenced by variation in micrometeorological conditions in different canopy crowns. Surprisingly, we did not observe significant or even moderate ^13^C enrichment from leaves to small twigs at all canopy heights on the three sampling days, except for 17:00–23:00 on 29 August (Fig. 4), which is similar to the findings of Gessler *et al*. (2004) who did not observe enrichment. However, previous studies have frequently reported that *δ*^13^C of leaf WSOM is more negative than that of twig phloem (Scartazza *et al*. 2004; Gessler *et al*. 2007; Eglin *et al*., 2009; Bogelein *et al*., 2012). This observation could be explained by a time lag for twig phloem. The ^13^C-enriched materials are produced in leaves but are not transferred into the phloem, depending on the relative allocation of new assimilates to ^13^C-enriched (starch) or ^13^C-depleted (lipids, lignin) carbon (Gessler *et al*., 2007), which might explain why WSOM in leaves is slightly ^13^C-enriched compared to that in twig phloem.

Furthermore, the results presented here indicated that significant ^13^C enrichment occurred when the water-soluble carbon was exported from twigs to stems (*p*<0.05; mean difference of 2.32 ± 0.65‰) (Fig.4). One hypotheses to explain the discrepancy is ^13^C fractionation associated with respiration. Recent reports claimed that δ^13^C of dark-respired CO2 by woody tissues is more positive than that of respiratory substrates (Damesin *et al*., 2005; Gessler *et al*., 2007; Maunoury *et al*., 2007). Brandes *et al*. (2006) also found a ^13^C depletion of 2.3–5.2‰ in stem phloem WSOM compared with emitted CO2. Another study in three herbs showed that CO2 respired by shoots was highly ^13^C-enriched relative to the respiratory substrate (Klumpp et al. 2005). These studies consistently support that fractionation during respiration caused ^13^C depletion in stem substrates. Thus, fractionation during respiration poorly explains phloem ^13^C differences between twigs and stems. Another possible explanation for the ^13^C enrichment in stems compared with twigs is the long time residence time for phloem carbohydrate transportation from twigs to stems. Assuming phloem transport rate ranges between 0.5 and 1 h (Zimmermann and Braun, 1971), it would take 8–16 h for phloem to be exported from a twig to the stem because the trees in this study are about 8 m tall. Gessler *et al*. (2004) also determined that ^13^C signals of stem base phloem were significantly related to the average daily values of stomatal conductance two days prior. However, the analysis of daily mean values showed that all twig phloem is significantly ^13^C-depleted compared with stem phloem (Fig.5). Therefore, time lag does not explain the twig-stem difference. The third hypotheses is that ^13^C enrichment might be associated with phloem transport. Carbon transportation in the stem does not cause ^13^C fractionation because *δ*^13^C of phloem sap does not change significantly in the basipetal direction (Gessler *et al*., 2008), but the partial retrieval of carbon is responsible for the difference of *δ*^13^C between twigs and stems. According to the recently reviewed dynamic Münch mass flow model, 2/3 carbon is transported back into the sieve tubes during the process of releasing carbon exported from twigs to stems (Minchin and Thorpe, 1987; Van Bel, 2003). The *δ*^13^C of retrieved carbon could be changed. Moreover, allocation of carbon to lignin, carrying lighter ^13^C signals for wood formation, can partly explain the twig-stem difference.

### ^13^C depletion of below ground organs

We observed root phloem has carbon isotopic composition similar to the sink tissue (leaves), with mean difference of 0.57±0.41‰. While compared with *δ*^13^C of stem phloem WSOM, there was significant ^13^C-depletion in root phloem (with mean difference of 1.85±0.64‰) (Fig. 4). Similar results have been found in rice (Deng *et al*., 2017). This effect might be ascribed to different dark respiratory mechanisms of different plant organs. Current research found that root-respired CO2 was ^13^C depleted in contrast to root carbon (Badeck *et al*., 2005; Klumpp *et al*., 2005; Bathellier *et al*., 2009). This result indicated that dark respiratory processes will not cause significant or even moderate ^13^C-depletion in root phloem WSOM. Another possible explanation is root carbohydrates could be recycled and transported in the acropetal direction (Heizmann *et al*., 2001). During the process of carbon transport from roots to stems or shoots, *δ*^13^C of root phloem WSOM might be altered. Moreover, some researchers suggested that root carbohydrates could also leak into the soil in the form of root exudates (Uchida *et al*., 2010; Werth *et al*., 2010). According to Uchida *et al*. (2010), 10% of carbon from atmospheric CO_2_ is transferred into soil by plant photosynthesis and carbon transportation. Deng *et al*. (2017) also found that 3.8%–10% soil organic carbon was derived from root exudates during the rice flowering period, and the percentage decreased along with the growth stages of the rice. Thus, the processes of carbon transport from root into soil should not be neglected, so the ^13^C signal of root phloem WSOM could also be modified by the process. However, the latter two hypotheses can be species-specific, resulting in different carbon fractionation patterns in roots compared with the carbon source (stem phloem WSOM). For example, the findings of Gessler *et al*. (2008) suggested that *δ*^13^C of phloem sap organic matter in stems at different sampling positions were similar to that in roots, and Badeck *et al*. (2005) analysed more than 80 publications and concluded that below ground organs were usually ^13^C-enriched relative to above ground organs. We also call on a better description of the characteristics of these hypotheses and what percentage of each process is responsible for post-photosynthesis fractionation in different plant organs.

### Day-to-day variation of δ^13^C is tightly correlated with *M*_s_

At diel time scales, there were moderate relationships between δ^13^C of leaves and *M*_s_ or *VPD*, while the relationship between *δ*^13^C of leaves and *M*_s_ was less evident (Table 1). The results indicated that δ^13^C of leaves is influenced by *RH* and *VPD* at short time scales and less dependent on *M*s which changes very little compared with *RH* and *VPD* during a day-night cycle. In contrast, *δ*^13^C of phloem WSOM in heterotrophic tissues was not strongly or even moderately dependent on these environmental factors as a result of a time lag. When diel variation is smoothed and daily mean values are calculated, we observed that both *δ*^13^C of leaf and twig phloem WSOM were strongly dependent on *M*_s_, and there was no dependence between other environmental factors and *δ*^13^C of leaf or twig phloem WSOM (Table 2). One thing to note is that *δ*^13^C of twig phloem WSOM is a better indicator of *M*_s_ than that of leaf WSOM. Merchant *et al*. (2010) also found that soluble leaf carbon, amino acid, and nutrient concentrations in the leaf were less predictive of plant water status than *δ*^13^C of twig phloem WSOM, although *δ*^13^C of leaf sugars and soluble extracts mirrored the variation in *δ*^13^C of twig phloem WSOM. A time lag for twig phloem carbon, which is a mixture of various carbohydrates with different metabolic histories, is insufficient to override the ^13^C signal of newly synthesized carbohydrates (Merchant *et al*., 2010). Moreover, leaf WSOM is more heterogeneous consisting of various sugars, organic acids, and amino acids with different residence times (Brandes *et al*., 2006; Gessler *et al*., 2008), while twig phloem WSOM is relative simple and is primarily composed of sucrose. This difference might additionally lead to *δ*^13^C of leaf phloem WSOM having relatively small responses to *M*_s_ compared with that of twig phloem WSOM. Thus, we propose that *δ*^13^C of twig phloem WSOM could be used as proxy for plant water status.

## Acknowledgements

This study was supported by the National Natural Science Foundation of China (No.41430747), the National Science Fund for Distinguished Young Scholars (No.41401013), and the Beijing Municipal Education Commission (CEFF-PXM2017_014207_000043).

